# Profile of the *tprK* gene in primary syphilis patients based on next-generation sequencing

**DOI:** 10.1101/422162

**Authors:** Dan Liu, Man-Li Tong, Xi Luo, Li-Li Liu, Li-Rong Lin, Hui-Lin Zhang, Yong Lin, Jian-Jun Niu, Tian-Ci Yang

## Abstract

**Background:** The highly variable *tprK* gene of *Treponema pallidum* has been acknowledged to be the cause of persistent infection. Previous studies mainly focused on the heterogeneity in *tprK* in propagated strains using a clone-based Sanger approach. Few studies have investigated *tprK* directly from clinical samples using deep sequencing.

**Methods/Principal findings:** We conducted a comprehensive analysis of 14 primary syphilis clinical isolates of *T. pallidum* via next-generation sequencing to gain better insight into the profile of *tprK* in primary syphilis patients. Our results based on primary syphilis clinical samples showed that there was a mixture of distinct sequences within each V region of *tprK*. Except for the predominant sequence for each region as previously reported using the clone-based Sanger approach, there were many minor variants of all strains that were mainly observed at a frequency of 1-5%. Interestingly, the identified distinct sequences within the regions were variable in length and differed only by 3 bp or multiples of 3 bp. In addition, amino acid sequence consistency within each region was found between the 14 strains. Among the regions, the sequence IASDGGAIKH in V1 and the sequence DVGHKKENAANVNGTVGA in V4 showed a high stability of inter-strain redundancy.

**Conclusions:** The seven V regions of the *tprK* gene in primary syphilis infection demonstrated high diversity; they generally contained a high proportion sequence and numerous low-frequency minor variants, most of which are far below the detection limit of Sanger sequencing. The rampant variation in each region was regulated by a strict gene conversion mechanism that maintained the length difference to 3 bp or multiples of 3 bp. The highly stable sequence of inter-strain redundancy may indicate that the sequences play a critical role in *T. pallidum* virulence. These highly stable peptides are also likely to be potential targets for vaccine development.

**Author summary:** Variations in *tprK* have been acknowledged to be the major contributors to persistent *Treponema pallidum* infections. Previous studies were based on the clone-based Sanger approach, and most of them were performed in propagated strains using rabbits, which could not reflect the actual heterogeneous characteristics of *tprK in vivo*. In the present study, we employed next-generation sequencing (NGS) to explore the profile of *tprK* directly from 14 patients with primary syphilis. Our results showed a mixture of distinct sequences within each V region of *tprK* in these clinical samples. First, the length of identified distinct sequences within the region was variable, which differed by only 3 bp or multiples of 3 bp. Then, among the mixtures, a predominant sequence was usually observed for each region, and the remaining minor variants were mainly observed at a frequency of 1-5%. In addition, there was a scenario of amino acid sequence consistency within the regions between the 14 primary syphilis strains. The identification of the profile of *tprK* in the context of human primary syphilis infection contributes to further exploration of the pathogenesis of syphilis.

## Introduction

Syphilis, caused by *Treponema pallidum*, is an ancient sexually transmitted disease that dates back to the 15^th^ century and is a public health threat that cannot be neglected [1, 2]. The completion of the first whole genome sequencing of the Nichols strain of *T. pallidum* provides a wealth of information about the characteristics of this pathogen, and since then the sequence of other experimental treponemal strains has also been released [3-8]. These particular achievements have revealed slight variations between different strains in a small genome (∼1.1 Mb), and most of the genetic diversity occurs in six genomic regions, including a polymorphic multigene family encoding 12 paralogous proteins (*tpr A* through *tprL*), highlighting most likely a factor in the pathogenesis of *T. pallidum* [2, 6, 9].

Within the *tpr* family, the antigen-coding *tprK* has been found to be the direct target of the human immune response [10]. Several remarkable studies performed in the rabbit model have demonstrated that the *tprK* gene possesses high genetic diversity at both the intra- and inter-strain levels, and the genetic variation in *tprK* is localized to seven variable regions (V1-V7) flanked by highly conserved domains [11-13]. Theoretically, through gene conversion, variations in the V regions would generate millions of chimeric *tprK* variants, resulting in a constant alteration in the *T. pallidum* antigenic profile [14]. Therefore, the *tprK* gene is acknowledged to have a pivotal role in immune evasion and pathogen persistence [15-17].

Previous studies have been mainly based on the clone-based Sanger approach; when using this approach, one would inevitably encounter a bottleneck in clone selection where minor variants, especially at low frequencies, are lost; consequently, the complete mutation profile of *tprK* is not fully understood. In addition, few studies have explored how *tprK* diversifies in the context of human infection, thus reflecting the actual heterogeneous characteristics of *tprK in vivo*, with the exception of one recent publication that reported on whole genome sequencing directly from clinical samples of *T. pallidum* [18]. Research has shown that after being cultured in rabbits, the genes of *T. pallidum* mutate and cannot retain their naïve characteristics, let alone the *tprK* gene, which has rampant potential to vary [19].

In the present study, we seek to systematically reveal the profile of *tprK* in *T. pallidum* directly from patients with primary syphilis by employing next-generation sequencing (NGS), thus providing important insights into the understanding of the diversity of *tprK* directly from primary syphilis patients and contributing to further explorations of the mechanisms of long-term *T. pallidum* infection.

## Methods

### Ethics statement

Written consent was obtained with signatures from all patients in accordance with institutional guidelines prior to the study. The study was approved by the Ethics Committee of Zhongshan Hospital, Xiamen University, after a formal hearing and was in conformance with the Declaration of Helsinki.

### Sample collection

Swab samples were obtained from the skin lesions of 14 patients (X-1~14) with primary syphilis. The clinical diagnosis of syphilis was based on the US Centers for Disease Control and Prevention (CDC) [20] and the European CDC (ECDC) [21].

### Isolation of DNA

Treponemal DNA was extracted from the swab samples using the QIAamp DNA Mini Kit (Qiagen, Inc., Valencia, CA, USA) according to the manufacturer’s instructions, and careful precautions were implemented to avoid DNA cross-contamination between isolates [22]. Each sample was quantified by targeting *tp0574* through qPCR using a 96-well reaction plate with a ViiA 7 Real-Time PCR System (Applied Biosystems, USA). A standard curve was constructed using 10-fold serial dilutions of cloned plasmids (for *tp0574*) generated through TOPO TA technology (Invitrogen, Carlsbad, CA, USA) and transformation of DH5α *Escherichia coli* cells [23]. The DNA samples that tested positive were stored at −20°C for further processing.

### Segmented amplification of the *tprK* gene

First, the extracted DNA was directly used in the amplification of the *tprK* full open reading frame (ORF). The primers used for the amplification are listed in Table 1. For amplification, KOD FX Neo polymerase (Toyobo, Osaka, Japan) was used. The reaction mixture contained 25 μL of 2× PCR buffer, 0.4 mM deoxynucleoside triphosphates, 0.3 μM of each primer, 1 U of KOD FX Neo polymerase, and 5 μL of genomic DNA in a final volume of 50 μL. The cycling conditions were as follows: 94°C for 2 min, followed by 40 cycles of 98°C for 10 s, 60°C for 30 s, and 68°C for 30 s. Then, the amplicons were gel purified and stored at −20°C for further processing as the next segmented amplification template.

**Table 1.**
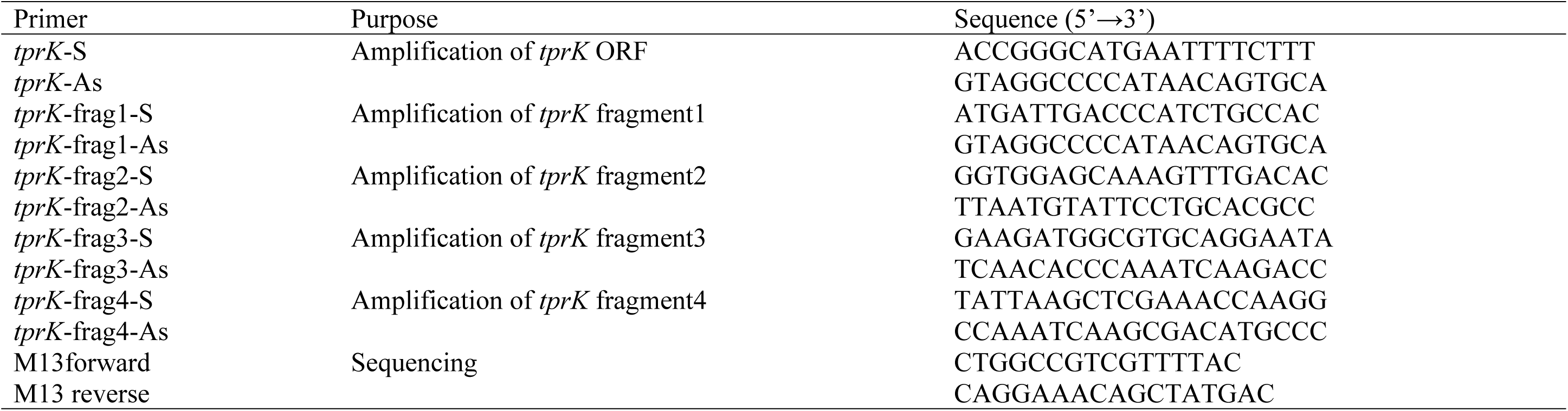
The primers for *tprK* amplification and sequencing.

Second, the *tprK* ORF was separated into four fragments overlapping by at least 20 bp (approximately 400~500 bp) for amplification. The primers are listed in Table 1. The purified DNA was diluted 1000 times and used as a template. The amplification mixture was the same as described above except that the primers were 0.15 μM. The cycling conditions were denaturation at 94°C for 2 min, followed by 30 cycles of 98°C for 10 s, 55°C for 30 s, and 68°C for 30 s. The size of all the products was verified by 2% agarose gel electrophoresis, and the products were gel purified. All purified amplicons were stored at −20°C for further processing.

### Library construction and next-generation sequencing

Library construction and sequencing were performed by the Sangon Biotech Company (Shanghai, China) on the MiSeq platform (Illumina, San Diego, CA, USA) in paired-end bi-directional sequencing (2×300 bp) mode. FastQC (http://www.bioinformatics.babraham.ac.uk/project/fatsqc/) and FASTX (http://hannonlab.cshl.edy/fastx_toolkit) tools were applied to check and improve the quality of the raw sequence data, respectively. The final reads collected from 14 patients were compared with the *tprK* of the Seattle Nichols strain (GenBank accession number AF194369.1) using Bowtie 2 (version 2.1.0).

An in-house Perl script was developed and applied to specifically capture DNA sequences within seven regions of 14 strains from raw data, both forward and reverse, as previously reported [18]. Briefly, the user-defined strings that matched the conserved sequence flanking the variable regions were used to catch the variable sequences. The defined strings referred to the mapping result of the reference and should be as long as necessary to ensure specificity (approximately 12-16 bp). Thus, the exact number of distinct sequences within seven regions across all strains was acquired. The intrastrain heterogeneous sequences were valid if the following conditions were simultaneously supported: 1) the number of the captured sequence was at least fifty reads and 2) the less frequent sequence displayed a frequency above 1%. Then, the relative frequency of the sequences within each variable region was calculated.

### *TprK* analysis by clone-based Sanger sequencing

An aliquot of DNA was also used for the amplification of the *tprK* full ORF according to the procedure described previously [11]. The purified amplicons were cloned into the pCR-2.1 TOPO vector (Invitrogen, Carlsbad, CA, USA) and were used to transform TOP10 competent *Escherichia coli* according to the manufacturer’s instructions. Approximately 10 clone plasmids from each sample were randomly selected and sequenced; each clone was sequenced not only in both directions with the M13 forward and reverse primers but also in the middle with the appropriate primers for a third reaction to ensure accuracy (Table 1). All sequencing was accomplished by the Bioray Biotechnology Company (Xiamen, China). The sequences within each intrastrain variable region were analysed using the BioEdit Sequence Alignment Editor Program (www.mbio.ncsu.edu/BioEdit/bioedit.html).

## Results

### 1. Description of clinical samples and *tprK* sequencing by NGS

The samples (N=14) were collected from patients diagnosed with primary syphilis at Zhongshan Hospital, Xiamen University. The clinical data of patients are shown in Table 2. The qPCR data of *Tp0574* showed that the number of treponemal copies in each clinical sample was eligible for the amplification of the *tprK* full ORF. The median sequencing depth of the *tprK* segment samples ranged from 10568.99 to 56676.38 and the coverage ranged from 99.34% to 99.61%, showing high homogeneity with the *tprK* gene of the Seattle Nichols strain.

**Table 2.** Description of clinical samples and *tprK* sequencing by NGS.

| Isolate | Gender | Age (year) | Serum RPR titer | Serum TPPA | Dark field microscopy | *T. pallidum* genome copies by *Tp0574* | Total reads | On-target reads (%) | Mean coverage of depth |
| --- | --- | --- | --- | --- | --- | --- | --- | --- | --- |
| X-1 | Male | 45 | 1:16 | + | Positive | 8.2E+03 | 357382 | 99.41 | 51967.28 |
| X-2 | Male | 27 | 1:16 | + | Positive | 8.82E+04 | 340240 | 99.47 | 49660.18 |
| X-3 | Male | 62 | 1:16 | + | Positive | 4.55E+04 | 398898 | 99.41 | 56676.38 |
| X-4 | Male | 65 | 1:4 | + | Positive | 1.15E+04 | 365060 | 99.34 | 52742.09 |
| X-5 | Male | 76 | 1:16 | + | Positive | 5.73E+04 | 363940 | 99.61 | 52960.83 |
| X-6 | Male | 64 | 1:32 | + | Positive | 2.33E+02 | 106934 | 99.37 | 14249.15 |
| X-7 | Female | 56 | 1:16 | + | Positive | 1.26E+04 | 114012 | 99.37 | 15579.12 |
| X-8 | Male | 46 | 1:4 | + | Positive | 1.41E+04 | 103280 | 99.43 | 12951.11 |
| X-9 | Male | 40 | 1:4 | + | Positive | 1.39E+03 | 119552 | 99.43 | 15864.28 |
| X-10 | Male | 66 | 1:32 | + | Positive | 9.17E+03 | 114064 | 99.37 | 14927.08 |
| X-11 | Male | 44 | 1:2 | + | Positive | 2.67E+02 | 94572 | 99.50 | 12935.89 |
| X-12 | Male | 39 | - | + | Positive | 6.40E+03 | 114588 | 99.43 | 14944.66 |
| X-13 | Male | 63 | 1:16 | + | Positive | 2.02E+02 | 118634 | 99.37 | 15013.54 |
| X-14 | Male | 61 | 1:1 | + | Positive | 1.16E+03 | 82812 | 99.37 | 10568.99 |

Abbreviations: NGS, next generation sequencing; RPR, reactive plasma reagin; TPPA, *T. pallidum* particle agglutination; +, positive; -, negative.

### 2. Sequence variability of *tprK* directly from primary syphilis samples

#### The number and length variation of distinct sequences in seven regions

According to the strategy, we extracted sequences within seven V regions to evaluate the sequence variability of *tprK* directly from primary syphilis samples. Altogether, 335 distinct nucleotide sequences were captured. The number of distinct sequences in the seven regions ranged from 21-76, with the highest number in V6 and the lowest in V1 across all samples (Fig 1). The length of the captured sequences within each region was also found to be variable, particularly in V3, V6 and V7, with 11 or 12 forms. In contrast, the length of the sequence in V5 had only two forms, namely, 84 bp and 90 bp. When the length of all sequences within each sample was calculated, the length of all distinct sequences differed by 3 bp or multiples of 3 bp. Interestingly, although the lengths of V3, V6 and V7 were particularly variable across all populations, these lengths continued to change by 3 bp. In this regard, the lengths of V1, V4 and V5 appeared to vary in intervals of 6 bp.

**Fig 1.**
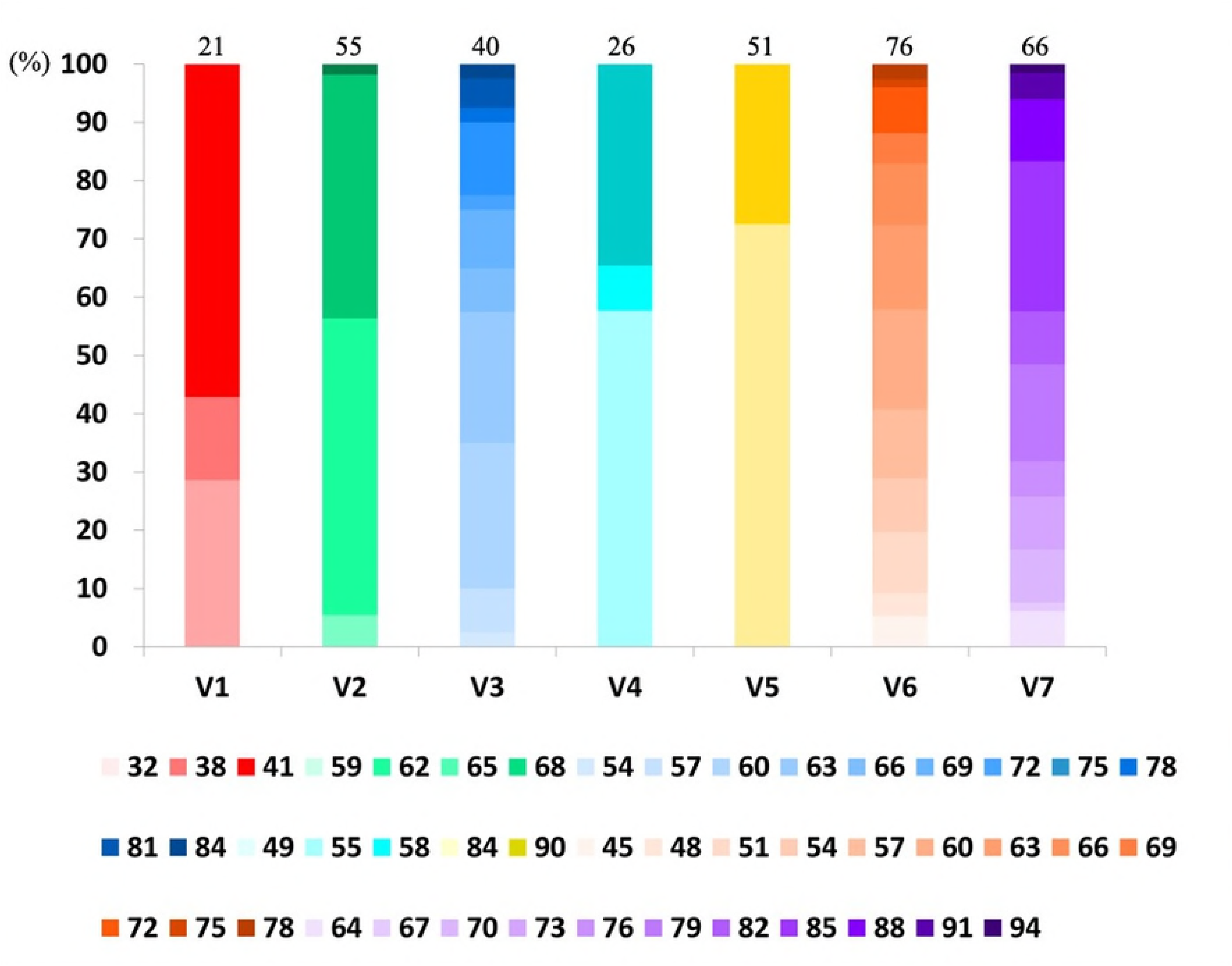
The varied length forms of distinct sequences within each region of *tprK*. The varied length forms within each V region are presented as the frequencies in each region and are filled with the gradient colour. All distinct sequences captured for each region are also shown above the V region.

#### The proportion distribution of distinct sequences in seven regions

The captured sequences were ranked by relative frequency within each V region of each strain. As Fig 2a shows, there was a predominant sequence in each region of all samples directly from primary syphilis patients, and the proportion of this sequence was almost 80%. It is worth noting that the frequency of the predominant sequence in some V regions of 4 samples (X-6, 8, 10, 13) was lower than 60%. In total, the frequency decrease appeared in the V2, V5, V6 and V7 regions, and the frequency in V6 of X-6 was even lower at 20.8%.

**Fig 2.**
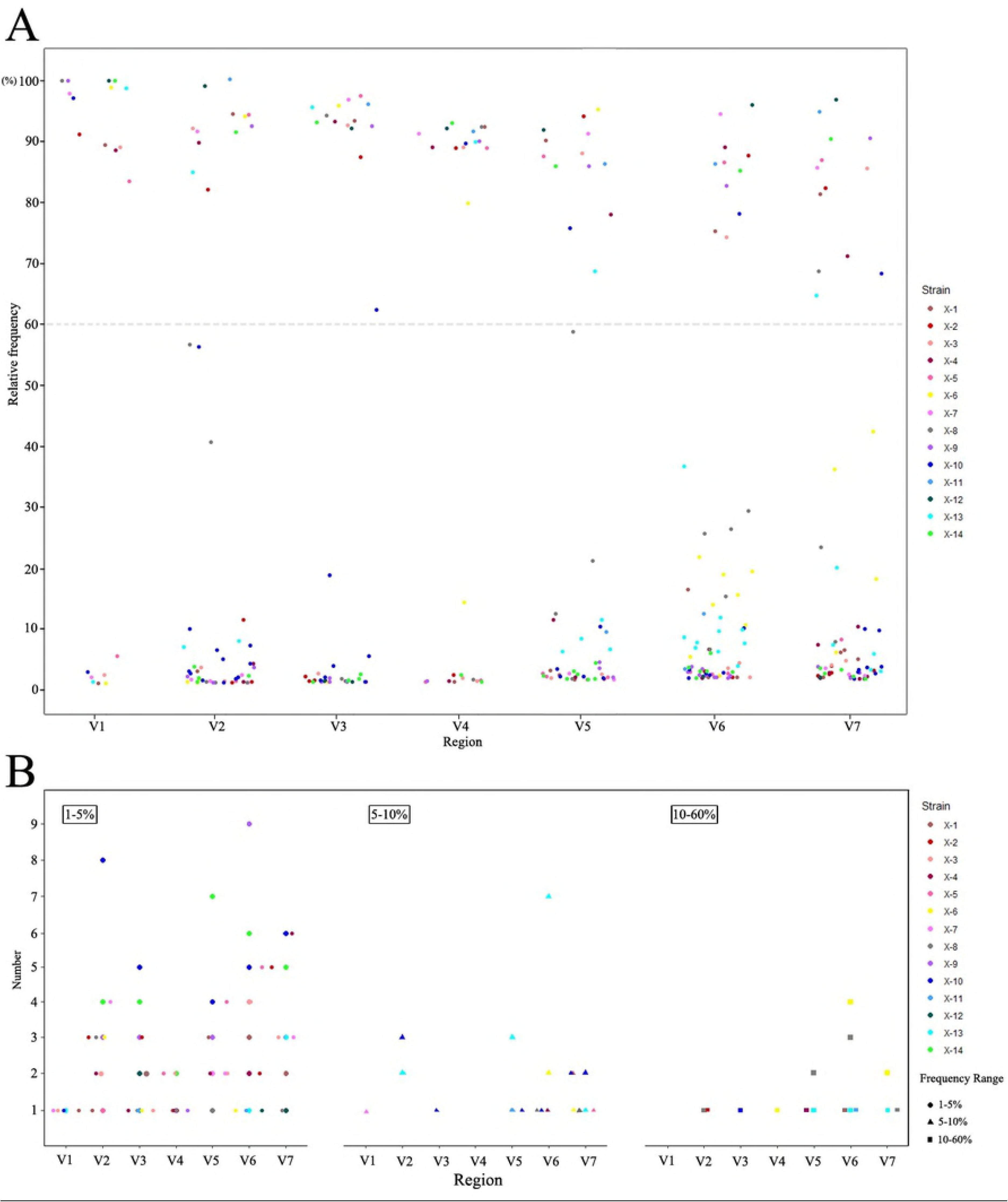
The proportion distribution of distinct sequences within each V region of *tprK*. (A) The dots indicate the relative frequency of identified distinct sequences within each V region of *tprK* across all 14 primary clinical samples, and the colour specifies the strain. (B) The graph shows the number of minor variants within each V region from all strains. The three thresholds (1-5%, 5-10% and 10-60%) are characterized by three different shapes, and the colour specifies the strain.

Apart from the detected predominant sequence within seven V regions, there was still a mixture of minor variants in each region. To investigate the relative frequency distribution of minor variants, we used three thresholds to explore the characteristics (Fig 2b). The major proportion of the variants in primary syphilis samples was in the 1-5% (181/237) range, and the lowest was in the 10-60% (22/237) range. At the two thresholds (5-10% and 10-60%), the observed variants were all mainly in V2, V5, V6 and V7 and from 4 samples (X-6, 8, 10, 13). This corresponded to the lower proportion of their predominant sequences.

### 3. Comparison with the heterogeneity of the clones within the population

Because previous studies were mainly based on the clone-based Sanger approach, we also applied this approach to analyse the *tprK* gene in these 14 strains and then compared the results with those of the NGS. Ten clones of each sample were obtained and identified by Sanger sequencing. Among the ten sequences, the predominant sequence within each V region of the primary syphilis samples was observed. The observed predominant sequence was consistent with the sequences obtained by NGS, such as in the strain of X-2 (Fig 3). However, where the frequency of the predominant sequence declined, especially when the frequency was less than 30%, for example, in the case of V6 in X-8, it became too ambiguous to distinguish the predominant sequences (S1 Fig). In all clones, the sequence was nearly undetectable for the minor variants with a frequency of 1-5%.

**Fig 3.**
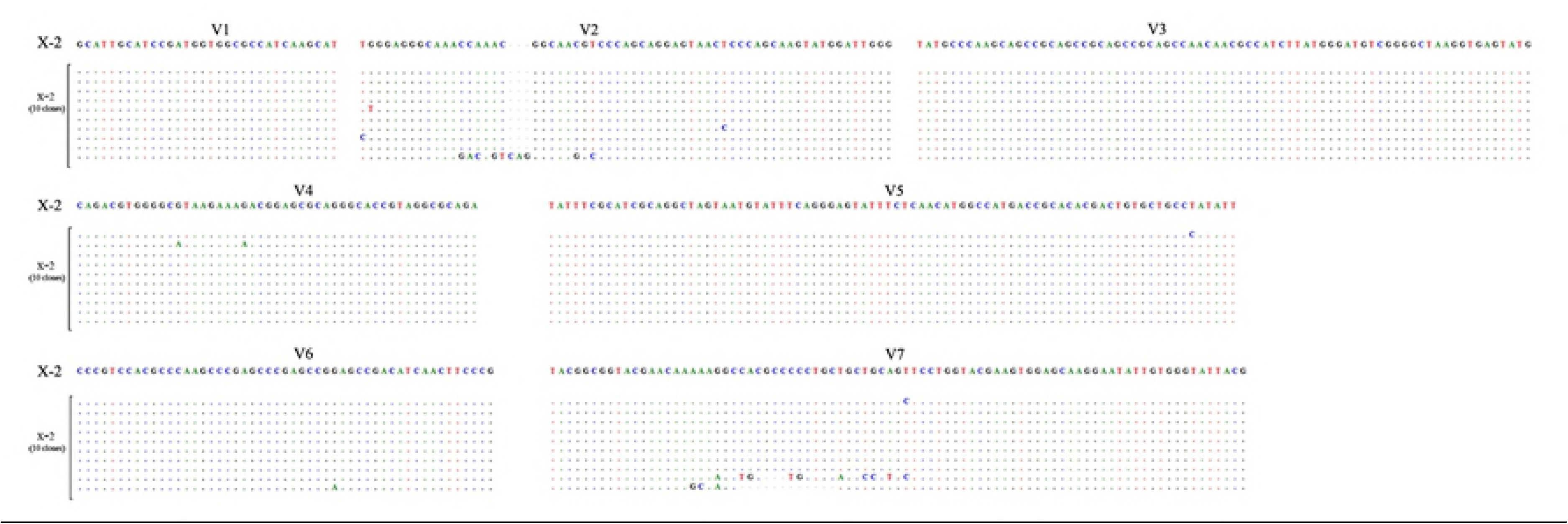
Predominant sequences within seven V regions identified by NGS compared to the results obtained by clone-based Sanger sequencing in this study. The alignment was performed on the X-2 strain as a representative sample. The identical nucleotides are shown on dots, and gaps in the sequence are shown by dashes.

### 4. Inter-population redundancy of the deducted amino acid sequence

A total of 335 nucleotide sequences were translated into amino acid sequences *in silico*. Ten sequences (10/335) were found to be synonymous, and at least 325 unique amino acid sequences were obtained. Unexpectedly, no sequence yielded a *tprK* frame shift or premature termination. When distinct sequences within each V region of all strains were compared, a scenario of sequence consistency was found. As Fig 4 shows, V1 and V4 presented a strong shared sequence capacity. The sequence IASDGGAIKH in V1 was observed in five strains (5/14) and DVGHKKENAANVNGTVGA in V4 was shared across seven strains (7/14). However, the parallel sequences in V3 and V6 did not seem as significant as in other V regions, especially in V6.

**Fig 4.**
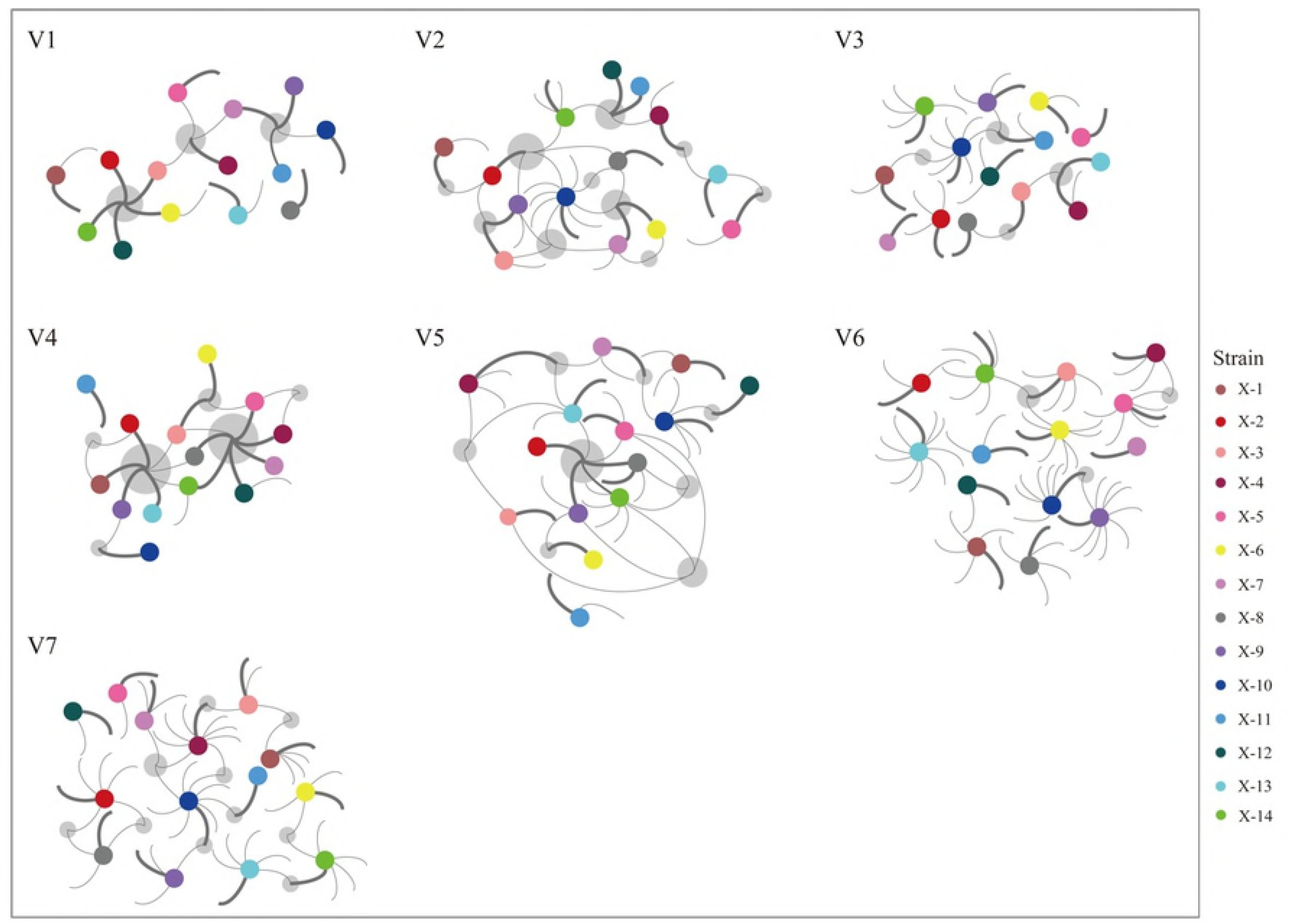
The scenario of redundant *tprK* amino acid sequences between all 14 primary syphilis clinical samples. The 14 strains are specified by coloured solid circles, and the predominant sequence and minor variants within each V region of one strain are represented by a bold arc and thin arcs, respectively. Each grey circle indicates the occurrence of sequence consistency between the strains.

To further explore whether the shared scenario was usually displayed by the predominant sequence across all the strains, we involved only the predominant sequence in the V region of each sample, which was represented by the bold arc in Fig 4, and found that V1 and V4 still presented similar shared sequence abilities despite the decreased redundant sequences. The occurrence of the consistent sequence in V1 and V4 could reach five strains and six strains, respectively. For the V3 and V6 regions, which were rarely consistent with sequences, the shared sequence in V3 occurred only between two strains, and there was no consistent sequence found in V6. Meanwhile, there was also no redundant sequence observed in V7.

## Discussion

Due to the inability to long term culture *T. pallidum in vitro*, research on this pathogen has been greatly hindered. The whole genome sequencing of the Nichols strain of *T. pallidum* provides a new perspective for the study of treponemal genes and proteins. Among these genes, *tprK* has been extensively studied because of its highly variable antigenic profile. It could effectively compromise the immunological function of specific antibodies generated by the host [14, 24-26] and cause immune evasion, thereby further leading to the development of late syphilis, neurosyphilis or serofast. Hence, intensive studies on the heterogeneity of *tprK,* especially in the context of human infection, would contribute to a deep understanding of the pathogenesis of syphilis.

In the present study, we performed NGS, a more sensitive and reliable approach, to gain better insight into the profile of *tprK* in primary syphilis patients. Overall, there was a wide sequence mixture focused on seven V regions of *tprK* in primary syphilis clinical samples. Among the seven V regions, V1 and V6 were known to have the lowest and highest variability, respectively [18, 27]. Our results also corroborated this feature in primary syphilis infection, in which the highest distinct number was found in V6 and the lowest distinct number was found in V1 (Figs 1 and 2a). Although *tprK* was revealed to have rampant genetic diversity within each strain [11, 12, 28], little is known about the exact proportion of these variant sequences within one strain. It is an advantage of NGS to fully discover the variants and determine the frequency [29, 30]. In this study, by using an in-house Perl script, we were able to retrieve the variants within the regions of each strain and calculate the relative frequency of the variants, thus disclosing the proportion of these variant sequences in primary syphilis patients. As shown in Fig 2a, there was a predominant sequence (the proportion above 80%) within each V region across all the strains.

In addition to the predominant sequence within each region, there was also a mixture of minor sequences (Fig 2a). Moreover, these minor variants were found to be mostly distributed at a frequency of 1-5% (Fig 2b), which was extremely below the detection limit for Sanger sequencing [31]. These results demonstrated that although the diversity of the V regions in antigen-coding *tprK* in primary syphilis infection was also presented to be wild, the V regions generally maintained their high proportion pathogenic sequence and numerous low-frequency minor variants. It is worth noting that the proportion of predominant sequences in some V regions of 4 samples (X-6, 8, 10, 13) was apparently lower than in others, and almost all the minor sequences in the 5-10% and 10-60% ranges were from these four samples, more specifically, mostly from the V2, V5, V6 and V7 regions (Fig 2). This result suggested that with the progression of disease or with increasing immunity, some V regions (V2, V5, V6 and V7) began to change. As a result, the frequency of the predominant sequence was decentralized. Instead, the frequency of a minor variant (or a new variant) gradually increased and further promoted the genetic diversity of *tprK* to escape immune clearance. Additionally, among the observed four regions, the frequency of the predominant sequence in V6 was particularly low (Fig 2a), suggesting that V6 may be the first affected region and is involved in immune evasion during the course of infection [14, 18].

We also applied the clone-based Sanger approach to analyse the *tprK* in primary syphilis patients in comparison with the results of NGS. As described in a previous study [32], the Sanger results generally displayed the predominant sequence within each region, which was consistent with the sequence found during NGS (Fig 3). However, for the lower frequency variants within the region, it became difficult to distinguish the predominant sequence, and we were unable to identify all the minor variants (S1 Fig). An increase in the number of clones selected would potentially alleviate this problem, but it would take more time and money. Additionally, the minor variants at a frequency of 1-5% were nearly undetectable in all selected clones. Therefore, use of the clone-based Sanger approach would lose much information about the complete profile of *tprK*, particularly in primary syphilis clinical samples in which *tprK* contained numerous low-frequency variants.

In this study, except for the distinct variations in *tprK* sequences, we found that the heterogeneity in *tprK* also presented in length (Fig 1). More length forms appeared in V3, V6 and V7, which was similar to the findings of Pinto et al. [18], demonstrating that the variations in these three regions could more easily cause changes in length. Despite this, the length forms were too far away to match with the number of sequences within each region; that is, the variants within each region were of the same length, but the context still had a considerable difference. For example, there were many different sequences observed in V5, but there were only two forms of length which was also observed in previous study [14]. This result indicated that the variants of *tprK* preferred a conversion without changing the initial length of the V region. Additionally, it was interesting that the length of all distinct sequences differed only by 3 bp or multiples of 3 bp, and previous research data also supported this pattern change [14, 18]. A multiple of 3 bp change pattern just matched with the triplet codon in protein coding, which has made us think about this feature probably explain why it is rare to uncover a *tprK* frame shift. It also suggests that the rampant diversity of *tprK* is regulated by a strict gene conversion mechanism.

Another noteworthy finding was the shared amino acid sequences across all the strains from the primary syphilis patients, which has also been observed in previous research [18, 27]. In our study, when all the distinct amino acid sequences within each region were aligned, at least half of the strains had sequences shared by other strains (Fig 4), which was similar to previous findings [18]. However, when only the predominant sequence within each region was analysed, *tprK* inter-population redundancy remained at a high level in only V1 and V4, in contrast to other regions, especially V6 and V7. This result suggested that the redundant sequences in V1 and V4 between strains were the ones that mostly dominated within a single strain. As the same antigenic sequences originated throughout the evolution of *T. pallidum* in different patients, this may reflect that the sequences become the best antigenic profiles to address the immune response of the host. The high stability of inter-population redundancy in V1 and V4 found in primary syphilis may confirm that the shared antigenic sequences in V1 and V4 play a critical role in *T. pallidum* virulence. In previous research [24, 25, 33], the molecular localization in the N-terminal region of *tprK* displayed promising partial protection in a rabbit model. Therefore, the highly stable shared peptide of V1 and V4 across all the strains would also likely be a potential target for vaccine development.

Finally, the limitations of our research should be discussed. First, the findings reported above lacked data about these clinical strains propagated by rabbits and could not directly highlight the difference from the naïve characteristics of *tprK* in human infection. This remains to be confirmed by animal experiments with NGS in the future. In addition, in this study, the number of clones selected for Sanger sequencing might be insufficient, although the current data were enough to verify the accuracy and reliability of our NGS results.

In summary, the characteristic profile of *tprK* in primary syphilis patients was unveiled to generally contain a high proportion sequence and many low-frequency minor variants within each V region. The variations in V regions were regulated by a strict gene conversion mechanism to keep the length differences to 3 bp or multiples of 3 bp. The findings could provide important information for further exploration of the role of *tprK* in immune evasion and persistent infection with syphilis. Furthermore, the peptide of each V region, especially the highly conserved peptide found in this study, could serve as a database of B cell epitopes of TprK for human immunological studies in the future.

## Supporting information

**S1 Fig. Comparison of the results of NGS and clone-based Sanger sequencing in V6 of the X-8 strain.** RF values indicate the relative frequency of each sequence.

S1 Table. The nucleotide sequences within the seven variable regions (V1-V7) of *tprK* captured directly from 14 primary syphilis clinical samples.

S2 Table. The amino acid sequences within the seven variable regions (V1-V7) of *tprK* captured directly from 14 primary syphilis clinical samples. * indicates synonymous nucleotide sequences within the same strain.

